# Whole-genome duplication and host genotype affect rhizosphere microbial communities

**DOI:** 10.1101/822726

**Authors:** Julian C. B. Ponsford, Charley J. Hubbard, Joshua G. Harrison, Lois Maignien, C. Alex Buerkle, Cynthia Weinig

**Affiliations:** Department of Botany, University of Wyoming, Laramie, WY, USA; Program in Ecology, University of Wyoming, Laramie, WY, USA; Marine Biological Laboratory, Josephine Bay Paul Center, Woods Hole, MA, USA; Laboratory of Microbiology of Extreme Environments, UMR 6197, Institut Européen de la Mer, Université de Bretagne Occidentale, Plouzane, France; Department of Molecular Biology, University of Wyoming, Laramie, WY, USA

**Keywords:** *Arabidopsis thaliana*, whole genome duplication, multinomial modeling, plant microbe interactions

## Abstract

The composition of complex microbial communities found in association with plants is influenced in part by host phenotype. Yet, the salient genetic architecture is often unknown. Genome duplication events are common in the evolutionary history of plants, influence many important plant traits, and may affect associated microbial communities. Using experimentally induced whole genome duplication (WGD), we tested the effect of WGD on rhizosphere bacterial communities in *Arabidopsis thaliana*. Specifically, we performed 16S rRNA amplicon sequencing to characterize differences between microbiomes associated with specific host genotypes (Columbia *vs*. Landsberg) and ploidy levels (diploid *vs*. tetraploid). We modeled abundances of individual bacterial taxa by utilizing a hierarchical Bayesian framework, based on the Dirichlet and multinomial distributions. We found that host genotype and host ploidy level affected rhizosphere community composition, for instance, the microbiome of the tetraploid Columbia genotype differed from that of other host genotypes. We then tested to what extent microbiomes derived from a given host genotype or ploidy level affected plant performance by inoculating sterile seedlings of each genotype with microbial communities harvested from a prior generation. We found a negative effect of the tetraploid Columbia microbiome on growth of all four plant genotypes. The findings suggest that while both host genotype and ploidy affect microbial community assembly, bacterial communities found in association with only some host genotypes may affect growth of subsequent plant generations.

**Importance:** Plants influence the composition of their associated microbial communities; yet the underlying host genetic factors are often unknown. Genome duplication events are common in the evolutionary history of plants and affect many plant traits, including the quality and quantity of compounds exuded into the root zone, which can affect root-bound microbes. In *Arabidopsis thaliana*, we characterized how whole-genome duplication affected the composition of rhizosphere bacterial communities, and how bacterial communities associated with two host plant genotypes and ploidy levels affected subsequent plant growth. We observed an interaction in which ploidy level within one host genotype affected both bacterial community composition and function. This research reveals how genome duplication, a widespread genetic feature of both wild and crop plant species, influences the coexistence of bacterial taxa and affects plant growth.

## Introduction

Plant-microbe interactions can exhibit a complete feedback cycle, in which changes in the microbiome affect plant performance and the genetics of the plant host alter microbial community composition (1–3). The traits of bacterial taxa in plant-associated communities affect both host fitness and ecological function (4). Rhizosphere bacteria, in particular, affect many aspects of plant performance, such as increasing access to nutrients (5), relieving abiotic and biotic stress (6), and promoting growth (7). Even slight changes in the rhizosphere microbiome can affect host plant performance (8). For instance, Korir *et al*. (2017) found that increased abundance of one taxon, *Bacillus megaterium*, led to enhanced nitrogen access and growth of *Phaseolus vulgaris* in field conditions (9). Wholesale changes in the abundance of taxa comprising rhizosphere microbiomes also affects plant performance (10, 11). For example, in *Arabidopsis thaliana*, differences in microbial community composition attributable to past or nearby plant communities strongly influenced host growth (12, 13).

A plant host’s genetic background can also affect the composition of microbiomes consisting of thousands of taxa (14–16). For instance, over a range of environmental conditions, host genotype in maize explains on average ∼19% of the variance in relative abundance of root microbial taxa (17). Among host genotypes, allelic variation segregating at loci with diverse functions could potentially contribute to differences in rhizosphere community composition. The contribution of specific host-plant loci and pathways has been demonstrated experimentally using genetic knock-outs or transgenic overexpression (18, 19). Genetic manipulation to shift the plant circadian clock by ± 4 hours explained ∼22% of the variance in rhizosphere bacterial communities among experimental *Arabidopsis* lines (13). However, the number of studies characterizing causal genetic factors is small, and the extent to which specific host genetic and genomic features alter the assembly and function of microbial communities remains largely unknown, despite the demonstrably important effect of the rhizosphere microbiome on host plant performance.

Plants influence rhizosphere microbial communities via root exudation of small molecular-weight organic compounds (20–22). For simple symbioses consisting of one plant and one microbial taxon, such as between legumes and *Rhizobia*, the host genetic mechanisms that underlie these interactions are increasingly well characterized (23–25). How more complex genetic mechanisms affect associated root microbial communities is less clear. Whole-genome duplication (WGD) is one genetic feature of host plants that could potentially influence microbial community composition via root exudation. WGD can lead to changes in cell size, life-history, and physiology, which may influence numerous other phenotypes (26, 27). WGD also occurs naturally in wild populations and is ubiquitous in the evolutionary history of plants and in the domestication history of many crop species (28).

In laboratory settings, colchicine is used to induce autopolyploidization and create stable lines of tetraploids in *A. thaliana*. This mutation theoretically produces no genetic changes besides those associated with genome duplication (29). Therefore, the comparison of rhizosphere microbiomes between *A. thaliana* tetraploid genotypes developed from inbred diploid lines provides a basis for testing the effects of host ploidy level on microbial community composition, without the confounding effects of allelic variation or fixed heterozygosity between tetraploids and diploid progenitors. Tetraploidy can induce changes in core metabolic pathways such as the tricarboxylic acid cycle (TCA), malate and citrate concentrations, and potassium uptake in *A. thaliana* (30, 31). These phenotypic shifts may alter root exudate profiles or other phenotypes that consequentially change microbial colonization and community composition.

In the current study, we were interested in characterizing the effects of host WGD on rhizosphere bacterial communities and testing how changes in bacterial community composition affected plant performance. We hypothesized that WGD would affect bacterial communities due to changes in host physiology induced by polyploidization. Further, if tetraploid plants assemble disparate communities compared to diploid hosts, then we predicted that differences in these communities could affect plant performance.

## Material and Methods

### Plant material and growth conditions

To test the effects of whole genome duplication and plant genotype on rhizosphere bacterial community composition and plant host performance, we selected two *A. thaliana* diploid genotypes: Columbia (Col-2x) and Landsberg *erecta* (Ler-2x) and their tetraploid counterparts (Col-4x CS922178) and (Ler-4x CS3900). Tetraploid genotypes were multiple generations removed from initial colchicine treatment, and therefore can be assumed to be mutationally stable (32). In our first experiment, each genotype was planted into sterilized potting mix and inoculated with a microbial community from the Catsburg region of Durham, North Carolina, USA (36.0622<sup>0</sup>N, -78.8496<sup>0</sup>W), a site with a history of *A. thaliana* growth. The microbiomes associated with each host plant genotype were characterized by 16S rRNA gene amplicon sequencing in the first experiment and tested for effects on growth of a second generation of plants in a second experiment.

In both experiments, seeds were surface sterilized using a solution of 15% bleach, 0.1% Tween, and autoclave sterilized reverse osmosis purified water (RO H_2_O). Seeds were thoroughly rinsed in RO H_2_O to remove any remaining detergents. Seeds were placed in 1.5mL tubes containing 1mL RO H_2_O and stored at 4°C for seven days and then placed on greenhouse benches under natural 14 hr photoperiods to induce synchronous germination (33). On the day root radicles were observed, seeds were transferred to 2-inch diameter net pots filled with a mixture of autoclaved Redi-Earth Potting Mix (Sungro Horticulture, Agawam, MA, USA) and 2 mL of liquid inoculant (described below). To ensure sterility, potting mix was autoclaved on a wet cycle for 60 min at 121°C, allowed to rest for at least one hour and autoclaved for another 60 min. No microbes could be cultured on tryptic soy agar media using serial dilutions of autoclaved soil as inoculum. To create inoculants for the first experiment, 60 g of the Catsburg soil was mixed with 540 mL of autoclaved RO H_2_O and sieved through 1000 µm, 212 µm and 45 µm sterile sieves to remove nematodes that could potentially affect plant performance (34). The Catsburg site not only has a well-documented occurrence of naturalized *A. thaliana* populations but also has a soil pH approximately identical to the potting substrate, which minimized potential selection by the common soil matrix on microbial community composition (35–37).

For the second experiment, we used four randomly selected plant-conditioned soil samples from each host genotype to create inoculums. Soil was combined by genotype, manually homogenized, and used to create inoculum as above. In all experiments, after ten days of growth, seedlings were thinned to one plant per pot. All experiments were performed at the Williams Conservatory or Agricultural Experiment Station at the University of Wyoming.

#### Experimental design

##### Experiment 1: Effects of host plant genotype and ploidy on microbial community

To measure the influence of host ploidy on rhizosphere bacterial community composition, 20 replicates of Col-2x, Col-4x, Ler-2x, and Ler-4x were planted in a fully randomized four block design. To avoid confounding the effects of ploidy with developmental stage, eight rhizosphere samples from each genotype were collected on day 24, by which time all plants displayed visible elongation of the primary inflorescence from the apical meristem (N = 32). Unharvested plants were allowed to grow until senescence to quantify additional traits. To account for variation in microbial communities due to greenhouse conditions and intrinsic variation in soil independent of plant host effects, we collected soil from empty pots containing only soil that were potted simultaneously with our experiment (N = 7).

##### Experiment 2: Microbial effects on plant performance

To investigate how host performance is affected by microbiomes associated with specific host genotypes, we used a fully factorial design with four host plant genotypes grown in four host-influenced microbiomes (harvested from the host genotypes in expt. 1). 40 replicates of each plant genotype were grown in sterilized potting mix inoculated with one of the four rhizosphere microbiomes from plant-conditioned soil in a fully randomized block design. Plants were checked daily for bolting and flowering. Upon the first plant bolting, a subset of plants was collected for above- and belowground biomass (N = 382) to measure plant growth and resource allocation. A subset of plants were allowed to senesce and seeds harvested to quantify seed mass (N = 152). All phenotypic measurements were performed following Rubin et al (38).

### DNA extraction and sequencing

Rhizosphere samples were collected from the root surface as described in (39) by first uprooting plants, manually agitating plants, removing loosely adhering soil particles, placing roots with closely adhering soil in a buffer of sterile PBS with 0.1% Silwet in a 15 mL tube, and vortexing for 10 min on maximum speed. Rhizosphere samples were centrifuged at 3714 rcf, and a total soil mass of no more than 250 mg was transferred from each tube into a sterile Qiagen PowerSoil (Qiagen, Valencia, CA) bead tube. DNA was extracted following manufacturer’s instructions. Extracted DNA was shipped to the Marine Biological Laboratories (Woods Hole, MA, USA) for amplification and sequencing of the V4-V5 (primers 518f and 926r) region of the 16S rRNA gene (40) on an Illumina MiSeq (Illumina, San Diego, CA, USA).

### Sequence and data analysis

We used the R package *dada2* (ver. 1.5.8) to filter and trim reads based on quality, estimate the error rate using 1 000 000 reads, dereplicate reads, infer amplicon sequence variants (ASV), merge paired end reads, remove chimeras, and assign unique sequences to taxa using the Silva 16S database (ver. 128) (41, 42). Next, we ascertained the relative abundance of operational taxa by quantifying unique sequences and amplicon sequence variants using the default settings in the *microbiome* R package (43). Raw sequences were uploaded to the NIH NCBI Short Read Archive under project ID PRJNA474006.

Dissimilarity analyses identify coarse-grained changes in community composition across treatments, which arise from the combined effects of all community members. Yet, significant differences in relative abundance may exist when comparing individual taxa between treatments that are not discernible when aggregating effects of whole communities. To test for differences in the relative abundances of individual taxa between treatment levels (contrasts of host ploidy and host genotype), we used a hierarchical Bayesian modelling approach reliant upon the Dirichlet and multinomial distributions (44, 45). The Dirichlet-multinomial model (DMM) explicitly accounts for the compositional nature of community sampling (whether in sequencing data or any finite number of taxa observations from an assemblage), readily allows information-sharing among replicates within a category and obtains estimates of the relative abundance of each microbial taxon, while propagating uncertainty in those estimates. We estimated differences in the relative abundance of each microbial taxon between experimental treatments through comparison of parameter estimates as per Harrison *et al*. (2020) (44).

Briefly, DMM estimates the multinomial parameters describing the relative abundances of each taxon in a replicate, denoted as a vector of parameters (***p***). These multinomial parameters were informed by a Dirichlet distribution, with parameters characterizing the expected frequencies of each taxon (**π\*** Θ), where **π** is a vector describing the expected relative abundance of each taxon in the sampling group and Θ is an intensity parameter that models among-replicate variation (44, 46). To quantify differences in the relative abundance of taxa between groups, we calculated the posterior probability distribution for the difference in π_i_ parameters between groups for some taxon *i*. The probability of an effect of treatment on a focal taxon can be determined by the location of zero in this distribution of differences. Following convention, if 95% or more of the distribution did not overlap zero, then there was little evidence that the relative abundance of the focal taxon differed between treatment groups.

The DMM offers several advantages over existing analytical methodologies. First, parameters were estimated while propagating uncertainty, thus avoiding cumbersome multiple comparison correction, and precluding the use of p-values. Second, information was shared among replicates and treatment groups. Third, rarefaction is unnecessary because analyses are performed on taxon frequencies, which are estimated using information from all replicates within a sampling group. Fourth, the Dirichlet and multinomial distributions have interdependent parameters that reflect the compositional nature of sequencing data, thus we model the composition as a whole as opposed to estimating the relative abundance of each taxon separately, as in DESeq2 (47).

DMM was specified in the Stan probabilistic programming language and implemented through the rstan (ver. 2.18.2) package in the R statistical computing environment (ver. 3.6.0) (48, 49). For each of four chains, the sampler was run for 1500 steps as a burn-in period and was followed by an additional 1000 steps (a total of 4000 samples were drawn from the posterior distributions of focal parameters [**π** from each sampling group]). The Gelman-Rubin statistic was computed to measure convergence among chains (50). Separate models were constructed for the following comparisons: all diploid versus all tetraploid hosts, Columbia versus Landsberg *erecta*, and Columbia tetraploid versus all other treatments (pooled).

For those taxa that differed credibly among our treatment groups, we used NCBI BLAST to confirm 16S rRNA identifications generated using the SILVA database (51, 52). We report species level identification in cases where reads align 100% with entries in the 16S ribosomal RNA sequences database, and report bacterial family if genus level resolution is not corroborated by SILVA (53).

To assess effects on plant performance in experiment 2, we used fixed effect, three-way ANOVAs, where inoculum, genotype, and block were the explanatory variables. For purposes of data representation, residuals were plotted after statistically accounting for the effect of genotype and block. Finally, we used planned comparisons to contrast plant performance between plants grown in the Col-2x versus Col-4x inoculums and Ler-2x versus Ler-4x inoculums.

## Results

### Sequencing results

After quality filtering and chimera removal in *dada2* as well as removal of bacterial taxa occurring in plant-less pots and removal of 16S rRNA sequences from chloroplasts, mitochondria, and Eukaryotes, 1 647 741 processed reads remained from an initial 3 282 410 raw reads. We retained 36 033 reads to 143 254 reads per plant rhizosphere (**Supplemental Figure 1)**. In total, there were 2689 microbial taxa (ASVs) present.

We utilized DMM to identify individual taxa with significantly different relative abundance due to host genotype and host ploidy level. This model allowed us to quantify differences in the abundance of individual taxa among treatments that might not be evident in whole community dissimilarity analyses. Based on this approach we identified 25 taxa that were significantly enriched in both Ler genotypes and 29 taxa that were more abundant in the Col genotypes (**Figure 1a)**. A taxon of genus *Pedobacter* was the most highly enriched in Ler associated bacterial communities with an 8-fold increase. In Col bacterial communities, the most enriched taxon was *Pseudomonas* at a 4.8-fold increase in abundance. DMM also identified 17 taxa that were significantly enriched in all diploid genotypes. Of these taxa, the most enriched taxon was a member of the genus *Bacillus*, enriched 3.1-fold relative to tetraploid communities. Tetraploid communities’ most enriched member was *Mucilaginibacter*, which was present at 4-fold greater relative abundance compared to rhizosphere microbiomes in diploids. 23 taxa were significantly enriched across both tetraploid genotypes (**Figure 1b)**. Our model identified 16 taxa as higher abundance in the Columbia tetraploid rhizosphere and 30 taxa enriched across the bacterial microbiomes of the other three host genotypes. In Col-4x, a bacterium identified as *Sphingomonas* had the highest change in abundance at 6-fold. Conversely, a taxon of genus *Pedobacter* was enriched in all other microbiomes. This taxon appeared at 5.8-fold greater frequency in these microbiomes than Col-4x. Across all comparisons, ASVs within the phylum Proteobacteria were more commonly differentially abundant than taxa within any other phylum.

**Figure 1:**
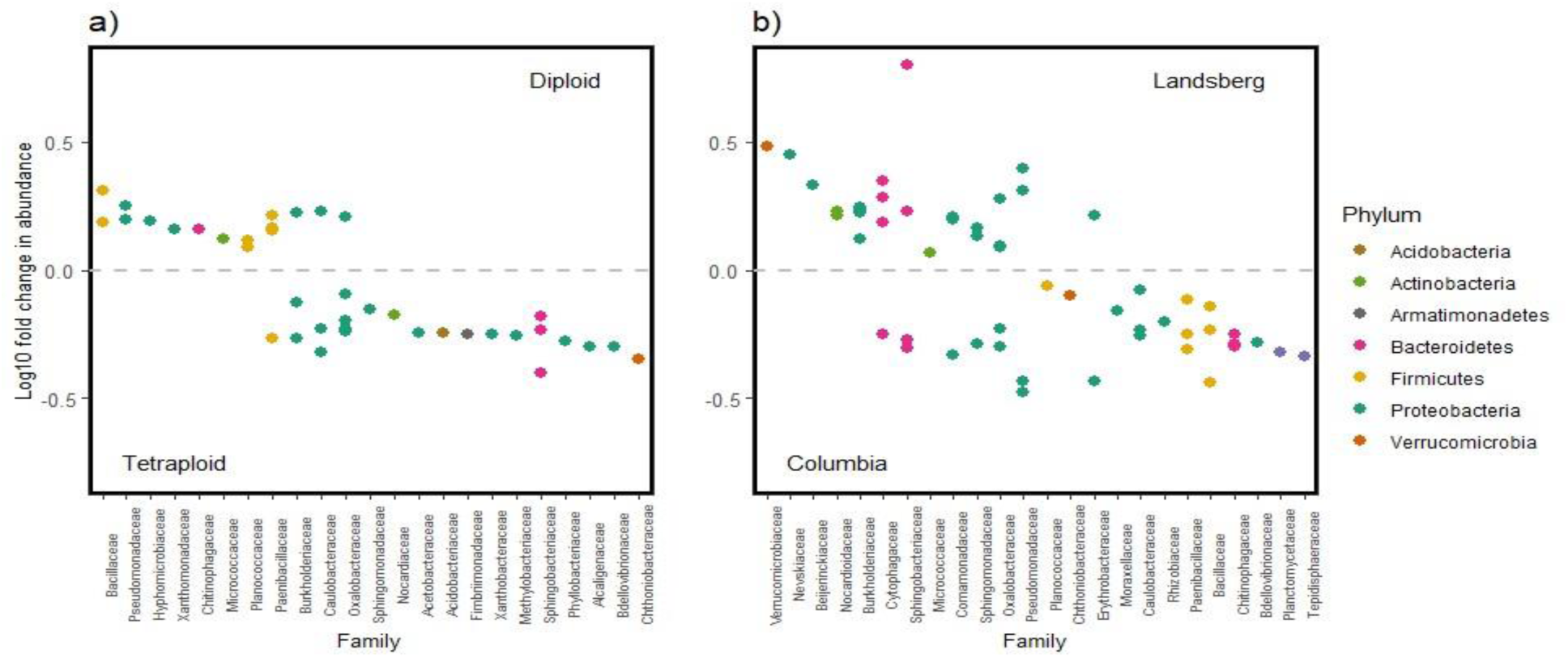
Genotype and ploidy differentially affect rhizosphere bacterial community abundances between the Col and Ler genotypes. a) Families identified as more abundant in the diploid rhizosphere are shown above grey line, and those more abundant in the tetraploid rhizosphere appear below grey line. b) Families identified as more abundant in the Landsberg rhizosphere are shown above grey line, and those more abundant in the rhizosphere of tetraploids appear below grey line. Log10 fold changes were calculated from relative abundance estimates obtained through hierarchical Bayesian modeling of read counts (see main text). Points represent individual ASVs within families.

### Effects of rhizosphere microbes on plant performance

When microbiomes from plant-conditioned soil were harvested following experiment 1, and used as inoculant in experiment 2, plants grown in soils inoculated with the Col-4x microbiome had significantly lower aboveground and belowground biomass compared to plants grown in soils inoculated with Col-2x microbiome (P < 0.001 & P = 0.012) or microbiomes influenced by either of the two Ler genotypes (**Figure 2ab**; P < 0.001 & P < 0.001). The soil microbiome had no effect on phenological characteristics or fruit number (**Supplemental Table 2**).

**Figure 2:**
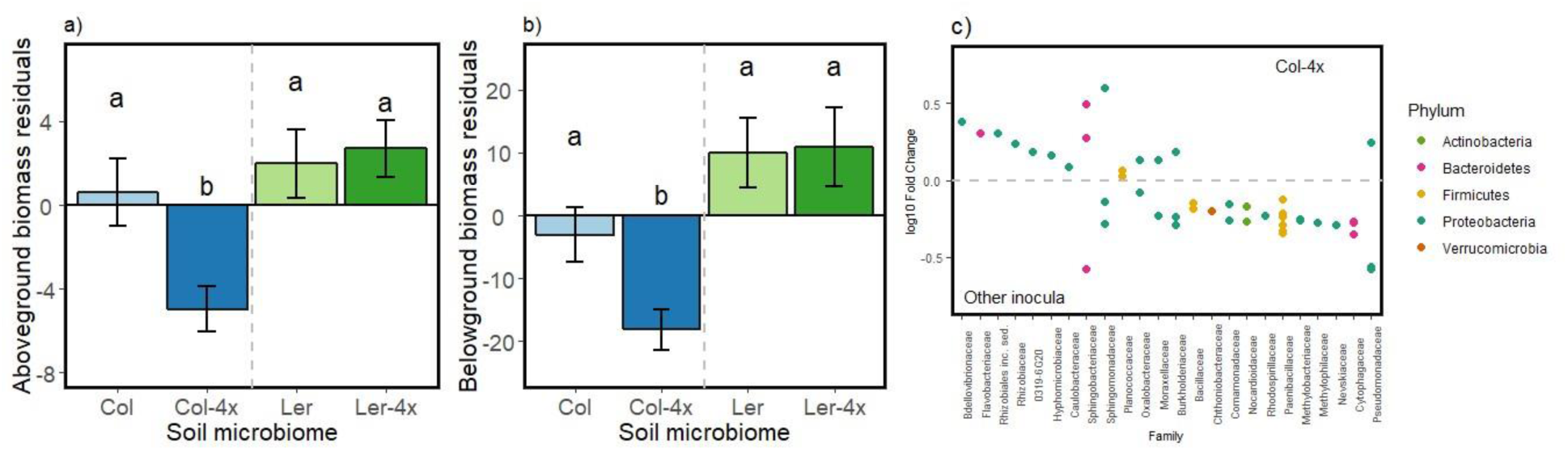
**a,b**) Residuals of observed growth differences in above and belowground biomass for plants grown in soils inoculated with microbiomes shaped by each genotype, with effects of block removed. All genotypes inoculated with Columbia tetraploid microbiome had significantly reduced above and below ground biomass. c) Bacterial families identified as more abundant in the Columbia tetraploid rhizosphere are shown above grey line. Those more abundant in the rhizosphere of all other inoculums appear below grey line. Log10 fold changes were calculated from relative abundance estimates obtained through hierarchical Bayesian modeling of read counts (see main text). Points represent individual ASVs within families

## Discussion

Whole genome duplication is estimated to have occurred across 30-70% of the angiosperm phylogeny over its evolutionary history and is a common genetic feature of many economically important plants (54). We found that host genotype and WGD influenced numerous individual taxa within the rhizosphere microbiome. Furthermore, host-genotype specific effects on rhizosphere microbiomes influenced plant growth. In contrast to prior studies (55), our experiments also utilized a soil with a history of *Arabidopsis* occurrence as inoculant. This should, in principle, allow genotypes to assemble root microbiota from the existing soil microbiomes.

Previous studies examining the root microbiota found in association with common lab accessions of *A. thaliana* have found a link between genotype and rhizosphere bacteria (15, 20). These studies report a < 10% shift in composition between the Ler and Col genotypes, which we corroborate here. As noted above, we identified many individual taxa associated with host ploidy level (**Figure 1a**). This provides support for the hypothesis that these specific taxa are directly responsive to genome duplication. Although the causal physiological mechanisms remain unclear in the current study, one candidate mechanism is root exudation. Previous studies suggest that carbon root exudates strongly affect the occurrence and abundance of microbial taxa in the rhizosphere (2, 56), and these exudates can be influenced by genome duplication (30). Consequently, shifts in root exudates, or other metabolites produced by the plant, in response to WGD could be partially responsible for the association between ploidy and the relative abundance of microbial taxa that we observed (57, 58). Given that the number of microbial taxa with a demonstrably significant response to WGD is in the 10’s rather than 100’s or 1000’s, we conclude that while host ploidy level affects the abundance of some taxa, it may not lead to broad-scale changes in the rhizosphere microbiome.

We found that differences in the rhizosphere bacterial microbiome lead to differences in host performance [**Figure 2**: 23, 61, 62]. Here, plants grown in soils inoculated with microbial rhizosphere communities harvested from Col-4x plants had reduced vigor compared to plants grown in soils inoculated with microbial communities from all other genotypes (**Figure 2**). One bacterial species, *Novaherbaspirillium autotrophicum*, was identified as being significantly enriched in the Columbia tetraploid genotype. *N. autotrophicum* was identified as a facultative autotroph capable of denitrification, isolated from rice paddy fields (61). The greater abundance of this microbe in the Columbia tetraploid community may point to changes in rhizobial community in plants inoculated with this microbiome. The greater abundance of *N. autotrophicum* may result in a decrease in available nitrogen in the rhizosphere causing lower overall growth in the plant host. This loss of biomass may have effects on nutrient cycling or survivability in stressful environments. Further, the effect of taxa such as *N. autotrophicum* may be especially impactful in the context of mixed populations of diploid and tetraploid individuals, where stressful conditions could be compounded or mitigated by host-associated microbes (62).

## Conclusions

In sum, we have shown that whole genome duplication influences rhizosphere bacterial community composition by affecting the relative abundance of certain bacterial taxa. In some genetic backgrounds, plant ploidy level affects microbial function, as indicated by enhanced above and belowground plant biomass in the Col-2x versus Col-4x genotypes. Multiple bacterial taxa are affected by host ploidy level, irrespective of host genetic background. The ASVs enriched (or reduced) in the rhizosphere of tetraploid hosts could help explain the advantages tetraploids appear to have in stressful environments (63, 64). Given the prevalence of WGD among plants in both natural and agricultural systems, our results highlight a novel mechanism by which plant evolutionary history influences the root-associated microbiome, thereby affecting plant phenotype.

## Acknowledgments

Computing was performed in the Teton Computing Environment at the Advanced Research Computing Center, University of Wyoming, Laramie (https://doi.org/10.15786/M2FY47). This work was supported by the National Science Foundation grants IOS-1444571 to CW, LM and EPS-16557226 to CW and CAB. We would like to thank Lindsay Leverett for collecting soil, and Meredith Pratt and Ryan Pendleton for help with experimental and greenhouse maintenance

## Conflict of Interests

The authors report no conflicts of interest.

## Supplemental Tables and Figures

**Supplemental Table 1:** Number of reads at each step of sequence data processing

| Processing Step | Number of Reads |
| --- | --- |
| Initial | 3 282 410 |
| Filtering and Trimming, Sample Inference,<br>Chimera Removal | 1 985 490 |
| Removal of Chloroplast, Mitochondrial,<br>Eukaryotic DNA and Plant-less blanks | 1 647 741 |

**Supplemental Table 2:** Effect of inoculum shaped by plant genotype on plant performance

| Trait | P - Value |
| --- | --- |
| Aboveground biomass | < 0.001 |
| Belowground biomass | < 0.001 |
| Root : Shoot | **ns** |
| Days to germination | **ns** |
| Days to flowering | **ns** |
| Seed Mass | **ns** |

## Notes

### Competing Interest Statement

The authors have declared no competing interest.

